# Single-cell Spatial Transcriptional Profiling Uncovers Heterogeneous Cellular Responses to Pathogenic Tau in a Mouse Model of Neurodegeneration

**DOI:** 10.1101/2025.11.19.689127

**Authors:** Xuehan Sun, Lidan Wu, Timothy C. Orr, Ashley Heck, Kimberly Young, Nathan Dunaway, Jose Ceja Navarro, Joseph M. Beechem, Miranda E. Orr

**Affiliations:** Department of Neurology, Washington University School of Medicine in St Louis, MO, USA; Bruker Spatial Biology, Inc., Seattle, WA; St Louis VA Medical Center, St Louis, MO

## Abstract

Intracellular tau accumulation contributes to neurodegeneration and systemic dysfunction. Although tau primarily aggregates in neurons as neurofibrillary tangles (NFTs), it also affects other cell types through poorly understood mechanisms. To define the molecular characteristics of tangle-bearing neurons and their influence on surrounding cells, we used CosMx SMI to evaluate the expression of 950 genes in more than 265,000 cells from rTg(tauP301L)4510 tauopathy and control mouse brains. In the cerebral cortex, tau pathology disrupted the excitatory-inhibitory neuron balance and altered pathways critical for myelination, similar myelin changes were observed in postmortem brain tissue from patients with progressive supranuclear palsy. In the hypothalamus, tau accumulation was associated with transcriptional changes linked to pathways involved in body weight and metabolic regulation. These findings reveal molecular mechanisms underlying both cell-autonomous and non-autonomous effects of tauopathy and identify targets for further investigation into tau pathogenesis and its systemic impact.

## Introduction

Tau was first identified as a microtubule-associated protein in 1975^1^. Although tau in its native form is highly soluble and shows little tendency to aggregate, pathogenic processes can lead to tau filament formation, driving a group of heterogeneous neurodegenerative diseases including, but not limited to, Alzheimer’s disease (AD), progressive supranuclear palsy (PSP), corticobasal degeneration (CBD), and some forms of frontotemporal dementia (FTD)^2,3^. In pathological conditions, tau proteins undergo numerous post-translational modifications, which impact their function and solubility, leading to aggregation and toxicity. Those toxic tau species, acting as seeding agents, can initiate the conformational change of endogenous tau in synaptically connected naïve neurons, thereby propagating the pathology^4–7^.

Neurons display different levels of vulnerability to tau pathology. For example, excitatory neurons from the entorhinal cortex are more susceptible to tau accumulation in AD. While some die early in the disease trajectory^8–10^, others undergo neurescence and survive long-term^11–13^. Whereas in PSP, tau aggregation starts in subcortical and brainstem neurons and glial cells, with subsequent spread to parietal and motor cortices, which associates with progressive movement impairment^14,15^. However, the mechanisms and molecular profiles underlying the selective vulnerability and resilience to pathogenic tau accumulation remain elusive.

Identifying tau-associated molecular changes with cell-type and brain-region specificity, while maintaining spatial resolution, is essential for understanding tau pathogenesis, yet been proven methodologically challenging. Experimental approaches to assess gene expression changes in response to pathogenic tau range from fluorescence *in situ* hybridization (FISH) and qPCR to measure single genes, to bulk tissue RNA sequencing and single-cell and single-nucleus RNA sequencing for a comprehensive assessment of the whole transcriptome. Collectively, these methods have revealed that many cell subpopulations undergo distinct transcriptional changes during disease progression, which may differ across brain regions, highlighting the heterogeneous and cell-specific response to AD pathologies^16–18^. Among these methods, only FISH-associated techniques preserve the spatial relationship between cells and tissue pathology, which is otherwise lost during tissue dissociation and cell isolation. However, the low-plex capabilities of FISH are not favorable when the experimental question requires information on hundreds or thousands of genes. To overcome this limitation, we used a spatial molecular imaging (SMI) technique that integrates single-cell transcriptomic with spatial information derived from brain pathology using in-tact histological brain sections^19^.

We employed CosMx SMI to characterize the transcriptomic changes, at subcellular resolution (∼50 nm), in the brains of rTg4510 tauopathy mice and non-transgenic littermate wild-type (WT) controls. This strain overexpresses human 0N4R tau carrying the P301L mutation (tau_P301L_) under the control of the Calcium/calmodulin-dependent protein kinase II (*Camk2a*) promoter that drives high levels of expression in forebrain neurons^20^. This model recapitulates key aspects of human tauopathy including neuronal loss, brain atrophy, and cognitive, physical and behavioral deficits^20,21^. We spatially mapped a curated list of 950 transcripts in brains from 20-month-old rTg4510 and WT mice, which reflect advanced disease stage comparable to the postmortem human brain tissues typically available in biorepositories. We customized the AT8 antibody, which detects phosphorylated tau, as a segmentation marker. This allowed us to directly visualize tau neuropathology across the tissue and perform molecular analyses on cells based on their spatial proximity to AT8-positive tangles. Differential expression (DE) analysis and the InSituCor method identified cell-type dependent and independent perturbations across different brain regions. These results provide new insights into the cellular and molecular changes associated with tau pathology at end-stage disease.

## Results

### Spatial molecular imaging analysis on rTg4510 mouse brain sections

Using the CosMx platform (Fig S1, see Methods for detailed description of the technology and workflow), we analyzed coronal brain sections covering hippocampal and adjacent regions from two 20- month-old rTg4510 and two aged-matched WT mice (approximately at Bregma position −1.955 mm). The sections were stained with AT8 antibody to detect tau phosphorylated at Ser202/Thr205, often used as a surrogate for NFTs. In rTg4510 samples, there was widespread AT8 staining throughout the cortex and hippocampus, and to a lesser extent, in the hypothalamus (Fig 1a). Antibodies against histone (nuclear) and GFAP (astrocytes), as well as an *in-situ* hybridization probe against 18S rRNA (cellular soma), were used to generate cell segmentation (Fig 1b).

**Figure 1.**
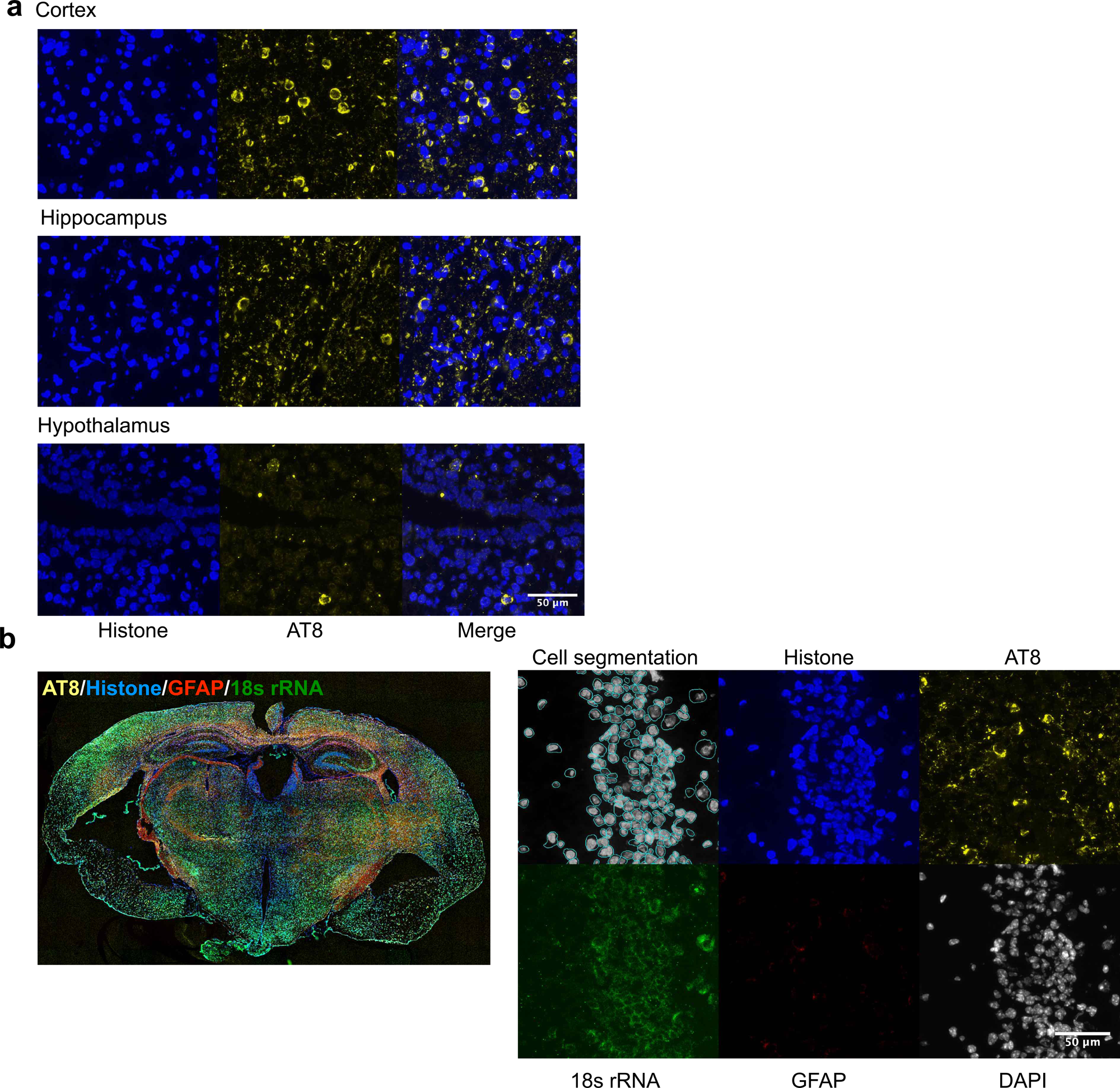
Mapping of transcription at single cell resolution using morphology-based cell segmentation by CosMx SMI. (a) Tangles were observed in cortex (top), hippocampus (middle), and hypothalamus (bottom) in rTg4510 brains. Tangles were stained by AT8 antibody (yellow). Nuclei were visualized by the antibody against histone (blue). (b) Representative images showing the segmentation markers in a coronal brain section of a 20- month-old rTg4510 mouse. Tau tangles were visualized by AT8 antibody (yellow). The other segmentation markers included antibodies against histone (blue), GFAP (red), and ISH probe against18s rRNA (green). Nuclei were counterstained with DAPI (gray). Segmented cell boundaries are outlined in blue.

We profiled a panel of 950 genes at single-cell resolution across the entire brain section. In this panel, 215 genes were selected for cell type identification, while the rest were curated to represent critical cellular processes such as neurotransmission, inflammation, and neuropathology (Fig S1d). Among the 267,072 imaged cells, over 98.91% of cells from each brain section passed quality control, and 788 out of the 950 panel genes were detected above background levels (Table S1). On average, 405 transcripts were identified per cell, with transcript counts ranging from 106 to 816 per segmented cell.

### Cell type Identification

We were able to identify expected major cell types using a supervised strategy (Fig 2a)^22^. Based on the original reference profiles^23^, 43 cell typing assignments were generated (Figs S2a and S2b), which were further collapsed into 10 major cell types (Fig S2c). To validate this classification, we also employed unsupervised cell typing that identified 11 major clusters (Fig 2b). Cell types were annotated based on specific gene markers (Fig 2c). For example, “neuron 1” were classified by high expression levels of genes related γ-aminobutyrate (GABA) production and storage, such as *Gad2* and *Slc32a1*, while “neuron 3a” and “neuron 3b” featured the enrichment of vesicular glutamate transporter *Slc17a7.* Other non-neuronal cell-type-specific markers included *Mog* for oligodendrocytes, *Vtn* for pericytes, *Csf1r* for microglia, and *Gja1* for astrocytes. Among these 11 cell types, oligodendrocyte a, neuron 1, and astrocyte were the most abundant, while oligodendrocyte precursor cells (OPC) were comparatively rare (Fig 2d). We found a high-level correlation between the two types of classifications as illustrated in the heatmap showing the number of cells that fell within each cell type generated by the two algorithms (Fig 2e). Therefore, both strategies detected similar major cell types, providing confidence in the cell typing, while offering slightly different resolutions at cell subtype level.

**Figure 2.**
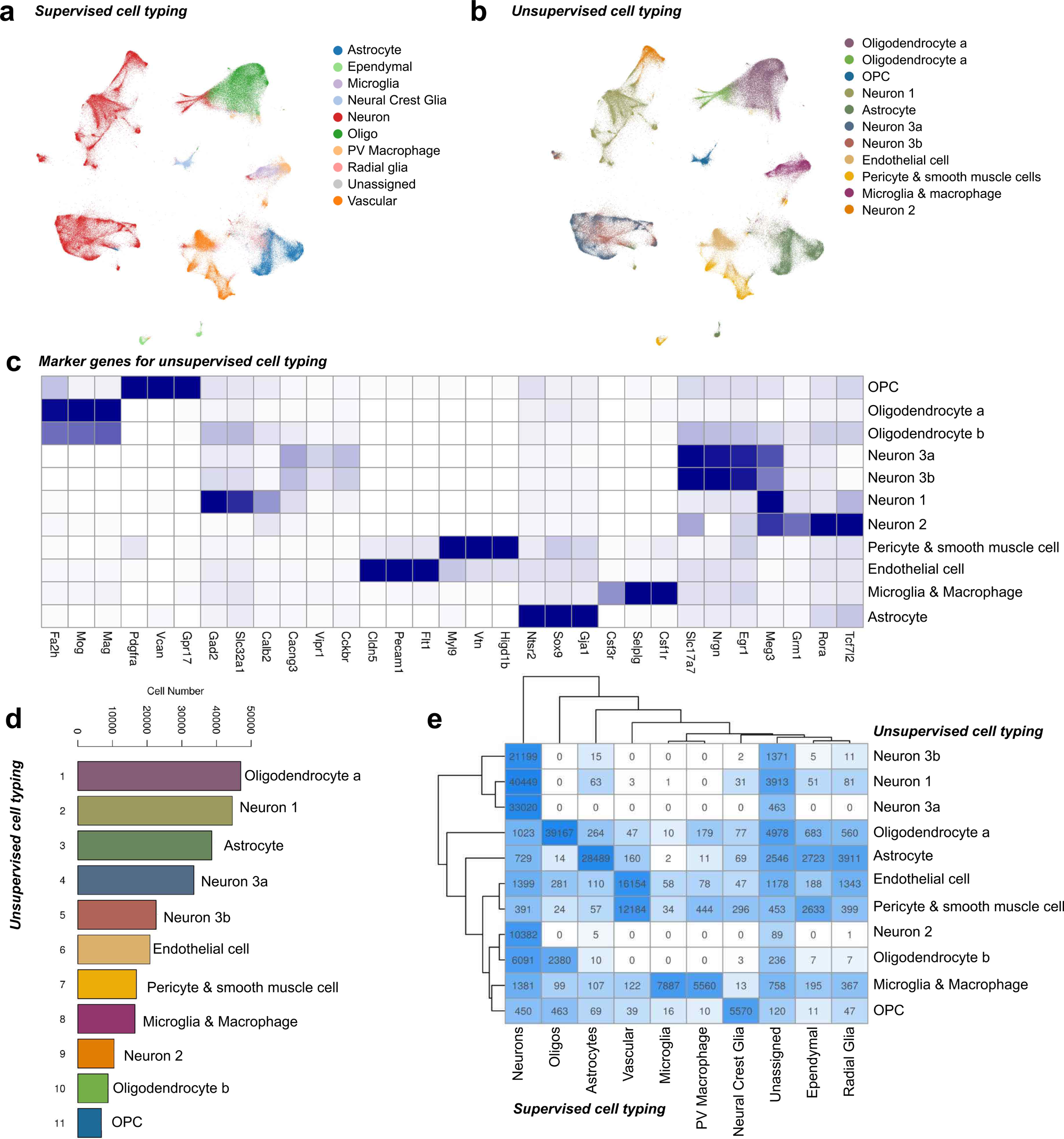
Cell type classification in the brain slices of rTg4510 and control mice. (a) UMAP plot showing the clustering of 267,072 segmented cells collected from the rTg4510 and WT samples (n = 2 per genotype). The negative binomial algorithm (supervised clustering) identifies 43 cell types based on a reference gene expression profile, which were condensed into higher levels and generated 10 major cell types (shown here). (b) UMAP plot showing 11 major cell types identified via unsupervised clustering. (c) List of marker genes used for cell type annotation after unsupervised clustering. (d) Statistics of unsupervised clustering. The data included all segmented cells from 4 samples. (e) Heatmap depicting the overlap in cell classifications between unsupervised and supervised clustering. The x-axis represents the 10 major cell types identified through supervised cell typing, while the y-axis corresponds to the 11 cell types derived from unsupervised clustering. The color intensity reflects the number of cells assigned to each intersection.

UMAPs of each sample revealed differential distributions within neuron clusters and the microglia/macrophage cluster between rTg4510 and WT mice, suggesting potential disease-associated changes at the cell type level (Fig 3a). Compared to the WT, rTg4510 samples showed a notable decrease of neurons (from 51.35% in WT to 34.75% in rTg4510, 38508 cells to 19749 cells), predominantly in the cerebral cortex. Conversely, perivascular (PV) macrophages (from 0.72% to 4.42, 537 cells to 2605 cells) showed an apparent increase in rTg4510 brains (Figs 3b and 3c).

**Figure 3.**
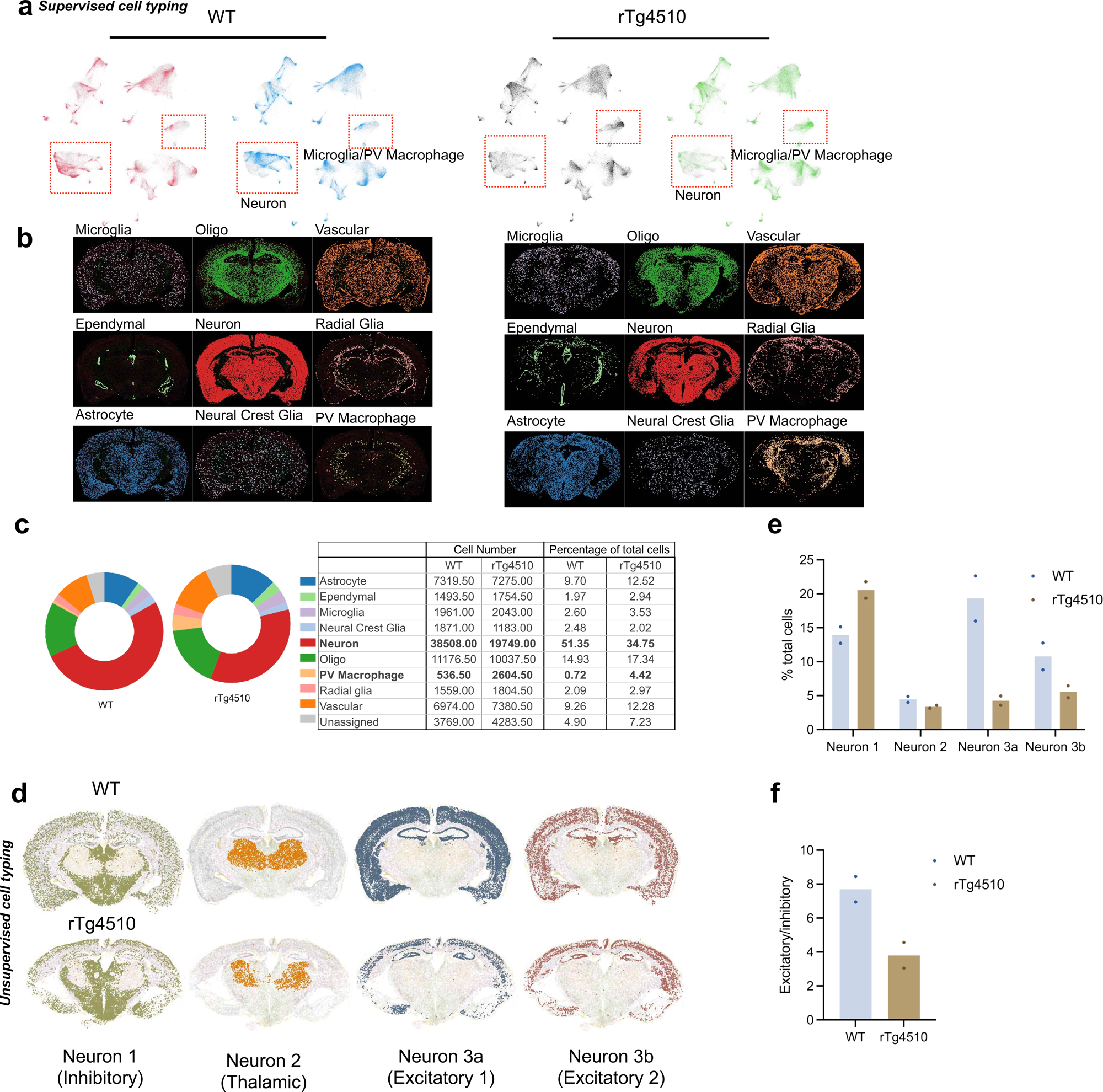
Cell type changes associated with tau pathology. (a) UMAP plots of individual samples. The differential clusters (neuron and microglia/PV macrophage) between 2 genotypes are highlighted in red boxes. (b) Representative spatial maps of each cell type in rTg4510 and WT brains. The cell typing was generated from supervised cell type classification. (c) Pie chart showing the cell type composition in rTg4510 and WT brains. The number and percentage of each major cell type were calculated as the average across biological replicates for each genotype. (d) Representative spatial mapping of neuron subtypes. The clustering was generated by unsupervised cell typing strategy. Cell type annotation in the parentheses was determined by their marker gene expression and regional location. (e) Bar plot showing the number of neuron subtypes identified by unsupervised cell typing. (f) Bar plot showing excitatory-inhibitory ratio in rTg4510 and WT brains (n = 2 per genotype). The calculation is based on excitatory neurons (3a and 3b) and inhibitory neurons (2), identified using the unsupervised cell typing strategy. Only neurons localized within spatial niches 1 and 2 (see Supplementary Figure 5) were included in the analysis.

More specifically, unsupervised clustering identified four neuron subtypes (Fig 2c). Based on their marker genes, neuron 3a and 3b were excitatory and neuron 1 was inhibitory. The neuron 2 cluster was also inhibitory, but expressed distinct transcriptional signatures including *Tcf7l2*, *Rora*, *Grm1*, and *Meg3*. The spatial cell maps showed that the excitatory neuron subtypes 3a and 3b were distributed throughout the cerebral cortex. Inhibitory neuron 1 cluster was present in both the hypothalamus and cerebral cortex, while inhibitory neuron 2 cluster was restricted to the thalamus (Fig 3d). We observed a decrease in the density of both excitatory neuron subtypes (3a: from 19.31% in WT to 4.26% in rTg4510; 3b: from 10.78% to 5.56%) and an increase in inhibitory neuron 1 in rTg4510 brains (from 13.93% to 20.56%) (Fig 3e). The ratio between excitatory and inhibitory neurons dropped from 7.70:1 in WT to 3.80:1 in rTg4510 in the cortical and hippocampal regions (Fig 3f).

### Susceptibility of hypothalamic cells to tau tangles

To better understand cellular responses to tau pathology at the molecular level, we first analyzed differentially expressed genes (DEGs) in neurons from rTg4510 and WT mice. We identified significant upregulated *Cartpt*, *Pomc*, and *Gal* (Fig 4a) in rTg4510 neurons, which encode neuropeptides regulating metabolic processes including food intake, energy expenditure, and body weight maintenance. The expression level of *Gal* and *Cartpt* showed a marked increase in the hypothalamus, where neurons and glia cells regulate energy homeostasis (Figs 4b and 4c). While *Camk2a*, the gene under the same promoter driving tau transgene expression, was largely restricted to the cerebral cortex (Fig S3a), we cannot entirely exclude the possibility of low-level transgene leakage in the hypothalamus. Therefore, the observed gene upregulation may be associated with either endogenous tau expression or pathology spreading to the hypothalamus.

**Figure 4.**
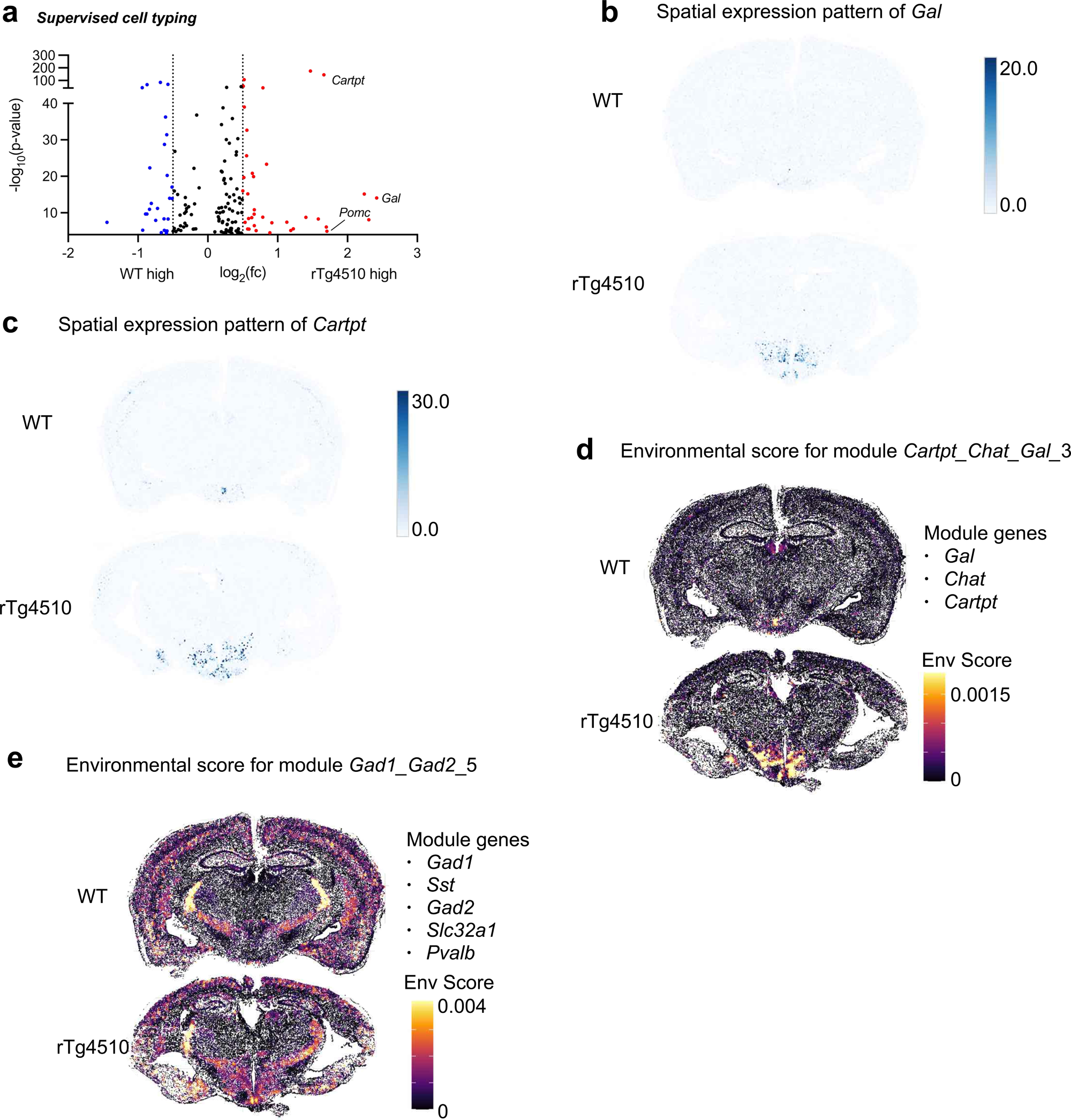
Differential expression and InSituCor analysis revealed the impact of tau pathology on thalamus and hypothalamus. (a) Volcano plot showing DEGs of neurons between rTg4510 and WT brains (y-axis: −log(p-value); x-axis: log_2_(fold change)). Selected genes (−log(p-value)> 4.3, |log_2_(fc)| > 0.5) are marked in red (up-regulated in rTg4510) or blue (down-regulated). Neurons were identified by supervised cell typing. (b) Representative spatial expression pattern of *Gal* in rTg4510 and WT brains. The shade of dots represents the number of transcripts detected in each segmented cell. (c) Representative spatial expression pattern of *Carpt* in rTg4510 and WT brains. The shade of dots represents the number of transcripts detected in each segmented cell. (d) Representative spatial overlay showing the environmental score for *Cartpt_Chat_Gal* module detected by InSituCor analysis. The environmental score was calculated as the average metagene score of each cell’s neighborhood, which summarized expression of genes belonging to the module. (e) Representative spatial overlay showing the environmental score for Gad1_Gad2_5 module detected by InSituCor toolkit. All 5 module genes were listed on the right.

To explore how cells relay the stress information from tau pathology to their surroundings, we performed InSituCor analysis^24^ that identifies gene co-expression modules independent from cell type landscape, and other confounding variables like sample-to-sample variance and background intensity. One module identified in this analysis consisted of 3 genes, *Gal*, *Chat*, and *Cartpt*. The environmental score map showed its peak in the hypothalamic region of the rTg4510 brain (Fig 4d). Another module of 5 genes also suggested the activation of inhibitory neurons primarily in the thalamus and hypothalamus in rTg4510 brains (Fig 4e). The gene content in this module implied an upregulation of GABA signaling (*Gad1*, *Gad2*, and *Slc32a1*) in somatostatin (*SST*) and parvalbumin (*Pvalb*)-positive interneurons. Loss of body mass commonly occurs during disease pathogenesis in rTg4510 and other tauopathy mouse models^25–28^, and was similarly observed in the mice included in our study (Fig S3b). Taken together, these data suggest that AT8-positive tau in the hypothalamus, either through tau spread or local production, as documented in mouse models and human patients^28–30^, coincides with the activation of pathways regulating energy intake and expenditure.

### Susceptibility of cortical and hippocampal neurons to tau tangles

We sought to characterize the neuronal transcriptional response to tau accumulation in the cortex and hippocampus. We first customized a “CorHipp” subtype to isolate cells in these 2 brain regions for DE analysis. Selection criteria included: (1) location within the regions of interest corresponding to cortex or hippocampus, and (2) assignment to niches 1 and 2 generated by single-cell neighborhood matrix, which were annotated as cortex and dentate gyrus, respectively. In WT samples, niche 1 captured cells in both the cortex and hippocampus. In rTg4510 samples, cells from these two brain regions clustered into separate niches. Niche 1 failed to capture cells in olfactory areas and most of the auditory areas in rTg4510 mice (Fig S4), likely due to the severe brain atrophy in this transgenic line at this disease stage.

Transcriptional changes in the CorHipp neurons highlighted synaptic function. Genes involved in calcium signaling, including *Camk2n1*, *Camk2d*, and *Calb1*, were downregulated in rTg4510 brains; whereas genes related to the complement system, including *C1qa*, *C1qb*, and *C1qc*, were upregulated (Fig 5a). Furthermore, the InsituCor analysis revealed one module of 33 genes (including *Snap25*, *Syt1*, *Syp*, and *Stxbp1*) that collectively indicated a decrease in synaptic transmission within the cerebral cortex of rTg4510 mice (Fig 5b).

**Figure 5.**
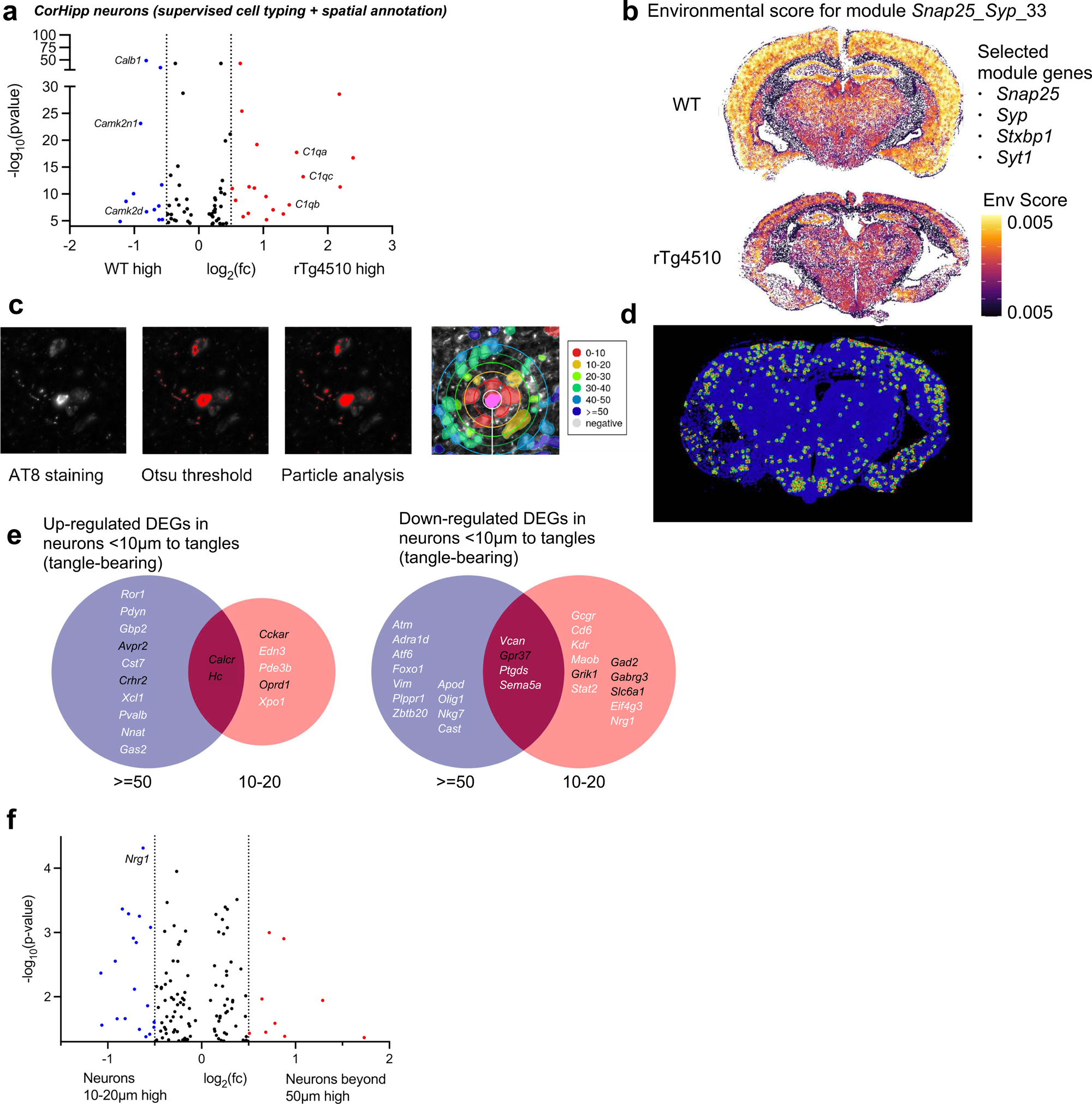
Cellular response to tangles. (a) Volcano plot showing DEGs of corHipp neurons between rTg4510 and WT brains (y-axis: −log(p-value); x-axis: log_2_(fold change)). Selected genes (−log(p-value)> 4.3, |log_2_(fc)| > 0.5) are marked in red (up-regulated in rTg4510) or blue (down-regulated). (b) Representative spatial overlay showing the environmental score for *Snap25_Syp_33* module detected by InSituCor analysis. 33 genes were detected in this module. Selected module genes related to synaptic transmission were listed on the right. (c) Identification of AT8 positive tangles. The AT8-positive stain was detected by Otsu auto thresholding algorithm, followed by particle analysis selection in Fiji. (d) A representative spatial map showing cells near segmented tangles in an rTg4510 brain section. Cells at various distances to AT8 positive area were highlighted with different colors. Cells far away from tangles (>= 50μm) were labeled in blue. (e) Venn diagram showing the DEGs from corHipp tangle bearing neurons compared to those in the proximal and distal environment. To ensure accurate estimation of backgrounds, all corHipp neurons were included in the analysis. DEG cut off: −log(p-value) > 1.3 and |log_2_(fc)| > 0.5. Genes of interest were highlighted in black. (f) Volcano plot showing the DEGs of corHipp neurons close to tangles (10-20 μm) to those far away (beyond 50 μm) (y-axis: −log(p-value); x-axis: log_2_ fold change). To ensure accurate estimation of backgrounds, all corHipp neurons were included in the analysis. DEGs (−log (p- value) > 1.3, |log_2_(fc)| > 0.5) are marked in red (up-regulated in tangle-bearing neurons) or blue (down-regulated).

Next, we assessed whether cells exhibited differential responses based on their proximity to tangles. Staining with the customized AT8 antibody allowed automated detection of tangle-bearing cells followed by segmentation to analyze their surrounding microenvironments. In rTg4510 mice, the AT8- positive area was detected by the Otsu auto thresholding algorithm in Fiji, followed by particle analysis selection with area ≥ 48.6 µm^2^ and circularity ≥ 0.27. No tangles were visible in the WT brains, and the cells were labeled as negative. Cells of interest were then grouped into six categories based on their distance from AT8-positive regions. The first five groups corresponded to 10µm intervals, representing distances progressively farther from AT8-positive cells, while the sixth group included cells located more than 50 µm away from NFTs (Figs 5c and 5d). Within the 0-10µm interval, we identified tangle- bearing neurons. Other nearby cell types and cells adjacent to potentially false-positive AT8 stains were also captured in this bin (Fig S5).

We focused the DE analysis on CorHipp neurons. Our DE model was designed to address potential imperfections in cell segmentation and batch effects of tissue sections, which improved accuracy by including more cells. Therefore, though we were mostly interested in the transcription profiles of tangle-bearing neurons (0-10µm) and those within their microenvironment (10-20µm), the model included all conditions within the CorHipp neuron subtype. To capture subtle differences between adjacent cells, we applied a less stringent p-value threshold in this analysis.

The 0-10µm bin capturing the tangle-bearing neurons, compared to their immediate neighbors, showed upregulation of *Hc*, a key component of the complement cascade, and downregulation of genes associated with synaptic transmission, including *Slc6a1*, *Grik1*, *Gad2*, and *Gabrg3*. Similar DEGs were observed when comparing these neurons to others located beyond 50 µm, suggesting tangle-specific effects. Interestingly, the bin with tangle-bearing neurons also upregulated receptors to various neuropeptides, including *Oprd1*, *Crhr2*, *Cckar*, and *Avpr2*, suggesting an enhanced responsiveness to neuropeptides. Non-tangle bearing neurons, regardless of spatial proximity to tangles, expressed higher *Gpr37*, a G-protein coupled receptor that regulates oligodendrocyte (OL) differentiation and myelination^31^, than NFT-bearing neurons (Fig 5e). Notably, neurons in the vicinity of tangles showed higher expression of *Nrg1*, a growth factor regulating myelination in both the peripheral and central nervous systems^32^, than neurons far from tangles (Fig 5f).

### Oligodendrocyte lineage alteration in the presence of tau tangles

Unsupervised clustering identified two OL subtypes and a group of oligodendrocyte precursor cells (OPC). Both subtypes expressed high levels of OL markers *Fa2h*, *Mog*, and *Mag* (Fig 2c). Cross- validation with supervised cell typing indicated that OL-a was primarily correlated with mature oligodendrocytes and myelin-forming oligodendrocytes (MOL and MFOL, respectively). In contrast, OL-b correlated with not only newly formed oligodendrocytes (NFOL) but also hindbrain excitatory neurons (HBGLU) (Fig S6).

Spatial cell maps and cell quantification revealed a notable increase in OL a density within the rTg4510 brains (Figs 6a and 6b). The cells exhibited a more expansive distribution throughout the cortical layers, compared to the relatively confined localization within the fiber tracts observed in WT brains (Fig 6a, left). Cell composition analysis showed an increase in OL a density (from 15.60% in WT to 20.25% in rTg4510), while the number of OPC remained relatively constant (WT: 2.71%; rTg4510: 2.33%) (Fig 6b). The statistics from supervised cell typing were in line with these results: while OPC numbers were comparable, MOL and MFOL density were higher in rTg4510 samples (Fig 6c, top). The data were then normalized against the total OL number, and we observed the same trend, suggesting that OPC differentiation and maturation patterns were different in rTg4510 mice (Fig 6c, bottom).

**Figure 6.**
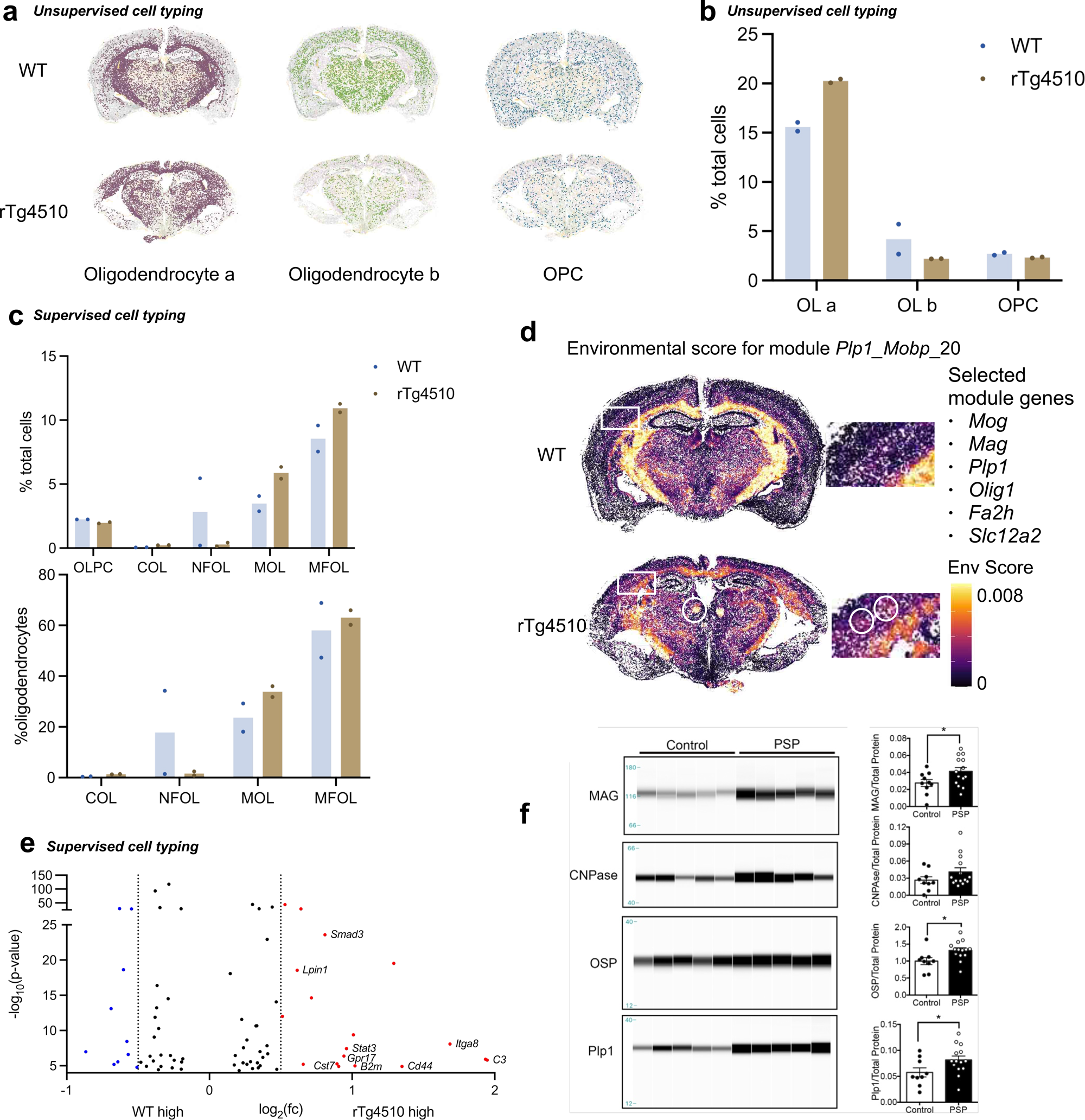
Tau pathology alters oligodendrocyte functions. (a) Representative spatial map of oligodendrocyte subtypes and OPC in rTg4510 and WT brains. The clustering was generated by unsupervised cell typing. Cell type annotation was determined by their marker gene expression. (b) Percentage of cells from each oligodendrocyte subtype and OPC identified by unsupervised Leiden clustering (n = 2 per genotype). (c) Cell type composition analysis based on supervised cell typing (n = 2 per genotype). Left: percentage of oligodendrocyte subtypes and OPC out of total number of cells. Right: relative enrichment of oligodendrocyte subtypes within the OL lineages. (OLPC, oligodendrocyte precursor cell; COL, committed oligodendrocyte; NFOL, newly formed oligodendrocyte; MOL, mature oligodendrocyte; MOL, myelin-forming oligodendrocyte). (d) Representative spatial overlay showing the environmental score for *Plp1_Mobp_20* module detected by InSituCor toolkit. 33 genes were detected in this module. Selected module genes related to myelination were listed on the right. (e) Volcano plot showing the DEGs of oligodendrocytes from rTg4510 to those from WT (y-axis: −log(p-value); x-axis: log2 fold change). DEGs (−log (p-value) > 4.3, |log2(fc)| > 0.5) are marked in red (up-regulated in rTg4510 oligodendrocytes) or blue (down-regulated). (f) Representative capillary Western blot analysis of myelin-associated proteins in temporal cortices from human PSP and ND controls. Quantification was performed on all samples (PSP: n = 14; ND: n = 10). Signals were normalized to total protein. Statistical significance was determined using Student’s *t*-test, and error bars indicate SEM.

InSituCor analysis revealed a change in distribution pattern of a module associated with oligodendrocyte maturation and myelination, which included key genes such as *Mog, Mag, Plp1, Olig1, Fa2h,* and *Slc12a2*. In rTg4510 brains, this module showed clustered upregulation in the cortical regions and thalamus, whereas in the WT samples, it was predominantly confined to the white matter (Fig. 6d). DE analysis between the two genotypes also revealed that rTg4510 OLs upregulated genes related to OL maturation and myelination, including *Smad3*^33^, *Lpin1*^34^, *Gpr17*^35^, and *Stat3*^36^. Additionally, genes involved in the immune response, including *C3*, *Itga8*, *Cd44*, *B2m*, and *Cst7*, were also upregulated (Fig 6e).

Lastly, we analyzed myelin protein expression in the temporal cortices of PSP patients, a human tauopathy also characterized by 4R tau aggregation. The mean total tangle score was higher in PSP patients (7.64) compared to non-demented (ND) controls (4.03; Table S2). Myelin-associated proteins, including myelin-associated glycoprotein (MAG), proteolipid protein 1 (PLP1), and oligodendrocyte- specific protein (OSP), were significantly upregulated in the grey matter from PSP temporal cortices compared to ND controls. However, levels of CNPase, a regulatory protein important for oligodendrocyte function and myelin maintenance, remained comparable between groups (Fig 6f).

Together, these findings suggest that tau pathology is associated with alterations in the oligodendrocyte lineage and myelin turnover in the brain.

## Discussion

We used high-resolution spatial transcriptomics (CosMx SMI), which detects RNA and protein within the same tissue section, to examine transcriptomic signatures in cells associated with tau pathology. Focusing on neurons with or without pTau-Ser^202^/Thr^205^ (NFTs) and cells in their microenvironment across cortical, hippocampal, and thalamic regions, we applied computational analyses which included cell typing, differential expression analysis, and spatial co-expression modules, to uncover widespread changes in cell composition in rTg4510 mice^37,38^.These analyses reveal imbalances in cortical excitatory/inhibitory neurons and altered thalamic neuropeptide signaling, consistent with seizure susceptibility and cachexia in human tauopathies. DEGs in neurons with or near NFTs highlight synaptic dysfunction linked to tau pathology. We also identified aberrant cortical myelination signatures in tauopathy mice, validated by elevated myelin protein expression in PSP patient tissues.

We quantified neuronal and glial cell composition changes in rTg4510 mice, which reflects global cellular changes reported in human tauopathy^37,38^ (Fig 3). We observed excitatory/inhibitory (E/I) imbalance in the cerebral cortex that potentially impairs neural circuit function and increases seizure susceptibility, which is more prevalent in patients with tauopathy than in other dementias^39–41^. While the loss of excitatory neurons may partially result from the *Camk2a* promoter driving tau^P301L^ expression in rTg4510 mice, tau itself—its expression level and genetic variants—modulates excitatory and inhibitory neurotransmission differently^42–44^. Tauopathy also alters neuronal synchrony, disrupts oscillatory activity, and heightens epileptogenesis risk in rTg4510 mice^45–47^. Although prior studies associate early tauopathy with hyperexcitability of excitatory neurons and increased seizure susceptibility, our data suggest that in advanced stages, shifts in E/I balance may also contribute to aberrant neuronal signaling.

Tau-induced neuronal loss has been attributed to excitotoxicity^48^, cell cycle re-entry^49,50^, and other apoptotic processes^51^. The decrease in high-quality cells due to pathology, along with imperfect cell segmentation, can confound DE analyses by introducing genotype-specific cell number imbalances. Therefore, we created a custom DE pipeline^52^, which employed negative binomial mixed model to identify DEGs while accounting for inherent subject-level and cell-level variability, and potential segmentation contamination. We identified downregulation of calcium signaling genes (e.g., *Camk2n1*, *Camk2d*, *Calb1*) in rTg4510 mice, consistent with reported disrupted calcium homeostasis in this and other tauopathy models^53^. In contrast, several genes related to the complement system were upregulated (*C1qa*, *C1qb*, and *C1qc*), implicating glial-mediated synapse elimination in neurodegeneration^54^ (Fig 5a). InsituCor analysis further revealed downregulation of a synaptic gene module (including *Snap25*, *Syt1*, *Syp*, *Stxbp1*) in rTg4510 cortices (Fig 5b). As our analysis was limited to a single timepoint, the onset of these changes during disease progression remains unclear. Investigating these phenotypes in early-stage tauopathy mice will be important in determining the time course of the phenotype and its association with neuron loss (e.g., upstream driver or downstream consequence).

NFT-bearing neurons, best characterized by the presence of phosphorylated, aggregated tau, remain poorly characterized transcriptionally. Prior works have used LCM or FACs sorting to collect neurons with and without NFTs^55,56^. Here we have identified a subset of DEGs in cortical and hippocampal neurons with NFTs (Fig 5). Notably, *Hc* (encoding C5 preproprotein) was upregulated (Fig 5e), linking NFTs to microglial activation and synaptic loss that accelerate AD progression^57,58^. These neurons also upregulated receptors to various neuropeptides, including *Oprd1* (to enkephalins), *Crhr2* (to corticotropin-releasing hormone), *Cckar* (to cholecystokinin), and *Avpr2* (arginine vasopressin) (Fig 5e), suggesting an enhanced responsiveness to neuropeptides. Interestingly, only neurons in the vicinity of tangles showed higher expression of *Nrg1*, a growth factor regulating myelination in both the peripheral and central nervous systems, suggesting a spatial-specific response to NFTs (Fig 5f). Additionally, non-tangle bearing neurons, regardless of spatial proximity to tangles, expressed higher *Gpr37*, a G-protein coupled receptor that regulates oligodendrocyte (OL) differentiation and myelination, than NFT-bearing neurons (Fig 5e).

Tauopathy mice consistently display age/disease-stage-associated weight loss compared to non- transgenic littermates^26–28^, whereas tau knockout mice display increased body mass^59,60^. We uncovered notable tauopathy-associated effects in the hypothalamus, and specifically in POMC/CART neurons that inhibit food intake and increase energy expenditure. The upregulation of *Cartpt*, *Pomc*, and *Gal*, could result from intracellular tau aggregation in the hypothalamus^61–63^. Alternatively, the upregulation could result from cell non-autonomous effects of pathology originating in distant brain regions, i.e., interfering with normal hypothalamus-hippocampal/cortical circuitry. Our results also revealed that tangle-bearing neurons upregulated receptors for various neuropeptides (Fig 5e). It is plausible that these neurons were responding to paracrine signals originating from these and other brain regions involved in integrating sensory, aversion, and satiety-related stimuli that modulate other aspects of feeding behavior^64^. Collectively, our data suggest that energy homeostasis dysregulation coincided with tau deposition in the hypothalamus.

The change of mature OL distribution coincided with a widespread spatial module responsible for myelination and axon ensheathment. Compared to WT mice, this module was downregulated in the white matter but upregulated in patches in the cortex of rTg4510 brains. This ectopic activation could result from localized axon injury. By analyzing postmortem human PSP brains, we observed elevated expression of proteins critical for compact myelin formation and axonal support in the grey matter from the temporal cortices, including in proteins encoded by the gene module shown to be ectopically expressed in rTg4510 gray matter. These data suggest oligodendrocyte dysfunction in rTg4510 brains featuring 4R tauopathy. Our DE analysis also showed that OLs in the rTg4510 mice upregulated genes involved in immune response. OL has been shown to adopt distinct transcriptional programs and cluster into different subgroups in response to tau and amyloid pathologies^65^. The proportional abundance in OLs we detected may represent the rise of disease-specific oligodendrocyte subpopulation driven by nearby tau accumulation or pro-inflammatory cytokines secreted by neighboring cells.

Collectively, our results highlight the broad impacts of tau pathology on transcriptional profiles among brain cells and across distinct brain regions. The integration of spatial information allowed us to reveal coordinated transcriptional responses among neighboring cells, suggesting that spatial proximity to NFTs may play a critical role in shaping cellular behaviors in the context of tau pathology, with an emphasis on oligodendroglia. Further exploration of this dataset, such as ligand-receptor analysis to elucidate direct communication between tangle-bearing neurons and neighboring cells, as well as trajectory analysis to model transcriptional transformations in different cell types based on their proximity to tangles^46,66^, will provide deeper molecular insights on cellular response to tau pathology.

## Limitations of the study

Our analyses provide important insights, while also highlighting areas for further investigation. For example, we used brain sections from discrete anatomical regions on a small number of biological replicates, which did not produce whole-brain data. We also focused on a single timepoint, end-stage disease, which precludes interpretations about disease trajectory. Moreover, our analysis utilized a panel of 950 curated genes, which likely did not capture a comprehensive assessment of all pathways participating in disease progression. To assess the translational relevance of tauopathy-induced oligodendrocyte changes, we included human brain samples. However, further studies in both experimental models and human tissue are needed to validate and extend these findings, including the cell subtype–specific transcriptomic signatures identified in this study.

## Resource availability

### Lead contact

Further information and requests for resources and reagents should be directed to and will be fulfilled by the lead contact, Miranda E. Orr.

### Materials availability

This study did not generate unique reagents.

### Data and code availability

- The CosMx raw data (metadata and associated count matrices) in this paper are available from the lead contact upon reasonable request.
- Code used for CosMx analysis is available from the lead contact upon request.

## Acknowledgements

M.E.O. is supported by the Alzheimer’s Drug Discovery Foundation (GC-201908-2019443), Cure Alzheimer’s Fund, Hevolution/American Federation for Aging Research, National Institute on Aging (R01AG068293, R01AG085182, R21AG087907, R01AG0909551, R25AG073119; U54AG079754, R01AG079224), National Institute of Neurological Disorders and Stroke (R56NS131387), the Rainwater Charitable Foundation, and US Department of Veterans Affairs (I01BX005717).

## Author contributions

Conceptualization: M.E.O., X.S., and L.W.; methodology: L.W., T.C.O., A.H., K.Y., N.D., and J.C.N.; analysis: X.S., L.W., and T.C.O.; writing—original draft: X.S. and M.E.O.; supervision: J.M.B. and M.E.O.; funding acquisition: M.E.O.

## Methods

### Mice

All animal experiments were carried out following the National Institutes of Health and Wake Forest School of Medicine Institutional Animal Care and Use Committee guidelines. We used 20-month-old female rTg4510 mice that reversibly express P301L mutant human tau on a Bl6/FVB genetic background. Non-transgene expressing littermates were used as controls. The founder mouse lines were acquired from the McLaughlin Research Institute for Biomedical Sciences, Great Falls, MT, and bred at Wake Forest School of Medicine, Winston-Salem, NC. Mice were anesthetized by isoflurane inhalation and euthanized by decapitation. One half was fixed in 4% PFA for 48 hrs and transferred to PBS/0.02% sodium azide at 4°C until further analysis. The other half was embedded in OCT, snap frozen in isopentane chilled with liquid nitrogen, and stored at –80°C until sectioning (5-um thickness) using a cryostat.

### Postmortem Human Brain Tissue

Postmortem human brain tissues (grey matter from temporal cortex) from PSP and ND patients were obtained from the Banner Sun Health Research Institute Brain and Body Donation Program (Sun City, Arizona). Detailed clinicopathological data, including postmortem interval, donor age and gender, as well as neuropathological assessments of amyloid deposition (stages A, B, and C) and neurofibrillary tangle progression (stages I–VI and total score), are summarized in Table S2.

### Tissue section preparation for SMI RNA profiling

Following the CosMx SMI RNA profiling protocol for fresh-frozen tissue sections (CosMx SMI Manual Slide Preparation User Manual, MAN-10159-03, Oct 2023 Ed, NanoString Technologies, Inc, Seattle, WA, USA. Available: https://university.nanostring.com/cosmx-smi-manual-slide-preparation-user-manual), we collected spatially resolved single-cell data from fresh-frozen, whole coronal brain sections containing hippocampus from two aged tau transgenic (rTg4510) mice and two age-matched WT controls. The RNA panel used is the CosMx Mouse Neuroscience panel, targeting 950 genes focused on robust neural and glial cell typing, neurodegeneration, and key aspects of cell state and signaling.

*In situ* hybridization on fresh-frozen tissue sections was performed using a procedure modified from existing protocols. Briefly, 5 µm-thick tissue sections were fixed for 2 h in 10% neutral-buffered formalin (NBF) at 4°C, then washed three times in 1X PBS for 5 min, once in 4% SDS for 2 min, and three times in 1X PBS for 5 min. After consecutive dehydration in 50% ethanol for 5 min, in 70% ethanol for 5 min, and in 100% ethanol for 5 min twice, sections were air dried for 30 min at room temperature. On the dried slides, incubation frames (Fream-Seal™ in situ PCR and hybridization slide chamber, Bio-Rad) were applied around the tissue sections. Proteins on the tissue were then digested with Protease IV (ACD Bio) spiked with Proteinase K (ThermoFisher) at 5 μg/ml at room temperature for 30 min. Tissue sections were washed twice with 1X PBS, incubated in 0.00015% fiducials (Bangs Laboratories) in 2× saline sodium citrate and Tween (0.001% Tween 20, Teknova) for 5 min at room temperature and washed with 1X PBS for 5 min. After protein digestion and fiducial placement, the tissue was fixed in NBF, washed in Tris-glycine buffer, washed in 1X PBS, blocked with *N*- succinimidyl (acetylthio) acetate (NHS-acetate, ThermoFisher), and washed in 2× saline sodium citrate (SSC) as described previously.

SMI ISH probes for panel targets and a morphology marker for the cellular soma, 16S rRNA, were denatured at 95 °C for 2 min and immediately cooled on ice. Probes were then added at 1 nM for panel targets and 0.2 nM for 16S rRNA to 40% formamide, 2.5% dextran sulfate, 0.2% BSA, 100 μg/ml salmon sperm DNA, 2× SSC, 0.1 U/μl SUPERase•In (ThermoFisher) in DEPC H_2_O. This ISH probe mix was added to the tissue, a seal was applied, and hybridization occurred at 37°C overnight. Following the ISH probe hybridization, tissue sections were washed twice in 50% formamide (VWR) in 2X SSC at 37 °C for 25 min and washed twice in 2× SSC for 2 min at room temperature.

Nuclear and morphology marker staining was performed to visualize cells for cell segmentation using the CosMx mouse neuron morphology kit which contains a cocktail of oligonucleotide-barcoded antibodies. Briefly, sections were stained with DAPI at 2 µg/ml in blocking buffer W (NanoString) to stain nuclei, washed in 1X PBS three times for 5 min, and then labeled with the barcoded antibody cocktail in blocking buffer W for 1 h at room temperature. The antibody cocktail contains an anti-histone H3 antibody for nuclei, anti-GFAP antibody for astrocytes, and anti-phospho-tau AT8 antibody for neurofibrillary tangle identification. Following antibody staining, sections were washed three times in 1X PBS for 5 min, blocked with 100 mM NHS-acetate in NHS-acetate buffer (0.1 M NaP + 0.1% Tween pH 8.0 in DEPC H_2_O) for 15 min at room temperature, and washed in 2X SSC for at least 5 min. The frame seal was removed, and a flow cell was attached to the slide for RNA profiling on the SMI instrument.

### Cell segmentation

The cell segmentation methodology used in this study integrates image preprocessing and machine learning techniques^19^. Briefly, tissues undergo staining with morphology markers, specifically DAPI and an anti-histone antibody for nuclear components as well as an anti-GFAP antibody and 16S rRNA-targeting hybridization probes for cellular soma. The cell segmentation pipeline takes those images and initiates rescaling, normalization, image deconvolution, and signal fusion processes, thereby producing dual-channel images representing nucleus-membrane features for each field of view. Utilizing pre-trained machine learning neural network models, Cellpose^67^, the pipeline conducts nuclear segmentation using the single nuclear channel and cytoplasm segmentation using the dual channels. Subsequently, the pipeline computes the Intersection over Union (IoU) scores between the two sets of segmentation results as the area ratio between the intersected region and the union of the two. The segments with IoU > 0.2 were merged, consolidating them into a singular cellular entity. This dual-mode approach is instrumental in accommodating staining non-uniformities. For the challenging task of capturing cellular protrusions in GFAP-stained images, the “Moments” auto-thresholding technique ^68^ is employed to delineate elongated cellular protrusions that may be difficult to capture with conventional machine learning methods. These protrusions are associated with their respective nuclei or soma using an intersection analysis between the protrusion masks and the outcomes of machine learning-based cell segmentation. This procedure enables the precise identification of the ultimate cellular boundaries.

### De novo cell clustering

Single-cell expression profiles were constructed by tallying the transcripts of each gene that were situated within the designated cell area as determined by the segmentation algorithm. Cells with fewer than 5 total transcripts were excluded from subsequent analyses. Data Normalization was performed by dividing each cell’s raw counts vector by its total counts. The Giotto package was then employed to process the normalized data through a series of steps for de novo cell clustering: dimension reduction using Principal Component Analysis (PCA), construction of Nearest Neighbor Network Construction based on the PCA outcomes, and Leiden Clustering on the resulting network. Subsequently, the identified clusters were further analyzed for marker gene identification and labeled with descriptive names given the marker genes. Giotto::runUMAP function was used to construct UMAP embedding based on sqrt-transformed normalized data, allowing the visualization of clustered cells in UMAP space.

### Supervised cell type annotations

The supervised cell type identification was performed with a likelihood-based clustering method, Insitutype^22^. Briefly, we derived cluster-mean profiles for each cell type using known cell type assignment and publicly available single-cell RNA-seq data for the mouse nervous system^23^. These cluster-mean profiles were then used as reference profiles by the Insitutype algorithm to identify anchor cells with the highest cosine similarity between their SMI raw expression profiles and the reference expression profiles of each cell type. The algorithm further updates the reference profiles by averaging SMI raw expression profiles within the identified anchor cells of the same cell type. Leveraging the anchor-derived reference profiles, Insitutype employs a Bayesian classifier to assign known cell types to individual cells. Cells with less than 80% posterior probability belonging to any known cell types were labeled as “unassigned”. In some analyses, the fine cell types identified by Insitutype were collapsed to major cell class level as indicated in Fig S2c.

### Identification of spatial interaction domains

Following the cell typing, we computed a single-cell neighborhood matrix that specifies the number of each cell type among each cell’s closest neighbors within an 80 µm radius in 2D physical space. The resulting matrix for each tissue section was processed independently and clustered using the Mclust algorithm to identify spatial interaction domains within each tissue section. For cross-sectional alignment, we carefully assessed the spatial domains concerning the mouse brain’s anatomical structure and the cell type composition. A neurologist assigned descriptive names to the spatially assessed domains, ensuring accurate alignment across multiple tissue sections, and associating these domains with specific brain structural features.

### Differential expression analysis

To mitigate the potential impact of cell segmentation errors in our differential expression (DE) analysis, we devised a customized pipeline^52^ that explicitly incorporates gene-level covariate for potential segmentation contamination in the DE model. Specifically, we initiated the process by employing the RANN package (https://github.com/jefferis/RANN) to identify neighboring cells within a fixed radius (50 µm in this study) but of different cell types from the cell of interest. We then calculated the total expression of each query gene in these neighboring cells of other cell types. This neighbor expression is weighted by (1/distance) of the neighbor, emphasizing the influence of closer neighbors. The resulting weighted neighbor expression of each cell was used to create a gene-level covariate (otherct_expr*_gene_*) that is useful for accounting for the potential spillover or segmentation error effects in the spatial transcriptomics dataset. In the downstream DE model, this covariate is transformed to standard normal distribution for stability, via a rank-based inverse normal transformation. To test whether each gene was differentially expressed between same cell types across different phenotypes (DE_variable), the negative binomial mixed models in nebula package (https://doi.org/10.1038/s42003-021-02146-6) were run predicting raw counts from the phenotype with the following formula, which also includes an offset term with log links using total counts of a given cell across all genes and a random effect term on different tissue sections.

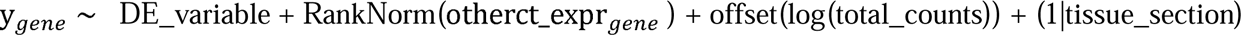

We employed the emmeans::contrast function (https://github.com/rvlenth/emmeans) to summarize the model predictions and performed the Wald test to test the difference in the expected expression rate between groups in the DE_variable (WT vs. rTg4510 in this study) for each gene.

### Spatial correlation analysis

We employed the InSituCor algorithm^24^ to investigate spatial correlations among genes within our dataset. Briefly, the algorithm first calculates the local gene expression environment of each query cell by averaging the raw gene expression across all neighboring cells within a 50-µm radius. To minimize the impact of unwanted variables like cell type, sample-to-sample variance, and background intensity, the algorithm constructs an “environment confounder matrix” summarizing these variables for each cell’s neighborhood. Subsequently, InSituCor derives the spatial correlation among genes by computing the correlation matrix of the environmental expression profiles while conditioning on the confounder matrix. The resulting conditional correlation matrix quantifies each gene’s propensity to be expressed in proximity of other genes beyond what can be explained by other confounding factors like cell type.

InSituCor further builds a gene correlation network based on the conditional correlation matrix. It identifies modules of genes that exhibit mutual co-expression patterns and conducts Leiden clustering to extract these modules. To aid in the interpretation and visualization of the spatial correlation results, InSituCor scores each identified gene module by computing a weighted average of its participant genes for both single-cell expression data and the cells’ environmental expression. InSituCor also summarizes the involvement of different cell types in each module, facilitating a comprehensive understanding of the spatial correlation patterns.

### Capillary electrophoresis immunoblots

To prepare the crude brain homogenate fraction, 50 mg of fresh-frozen brain tissue was homogenized using a Dounce homogenizer (20 strokes) in Cytoplasmic Extraction Buffer (CEB) from the Thermo Scientific Subcellular Protein Fractionation Kit for Tissues, supplemented with 1× Phosphatase Inhibitor Cocktail II and Halt Protease Inhibitor Cocktail (Thermo Scientific), at a 10% (w/v) ratio. A 100 µL aliquot was removed, subjected to sonication using a Fisher Scientific Sonic Dismembrator Ultrasonic Processor FB-120 (three 5-second pulses at 20% amplitude, with 30-second intervals between pulses), and clarified by centrifugation at 21,000 × g (14,800 rpm) for 20 minutes at 4°C. The supernatant was collected, and protein concentration was determined using a BCA assay. Lysates were diluted to 0.4 µg/µL in 0.1X Wes Buffer for capillary electrophoresis using the HRP-luminol/peroxide detection system. Primary antibodies were diluted in Wes Antibody Diluent as follows: CNPase (1:100, Cell Signal Technology, D83E10), MAG (1:200, Cell Signal Technology, D4G3), Plp1 (1:100, Sigma, HPA004128), and OSP (1:50, Abcam, ab53041). Signals were normalized to total protein.

## Acknowledgements

MEO is supported by the Alzheimer’s Drug Discovery Foundation (GC-201908-2019443), Cure Alzheimer’s Fund, Hevolution/American Federation for Aging Research, National Institute on Aging (R01AG068293), National Institute of Neurological Disorders and Stroke (R21NS125171), the Rainwater Charitable Foundation and US Department of Veterans Affairs (I01BX005717).

## Disclosures and completing interests

L.W., A.H., K.Y and J.M.B are employees of Bruker Spatial Biology, Inc. J.M.B. also is a shareholder of Bruker Spatial Biology, Inc. N.D and J.C.N were former employees of NanoString Technologies (now Bruker Spatial Biology). M.E.O. has a patent “Detecting and Treating Conditions Associated with Neuronal Senescence”.

## Figure Legend

**Supplementary figure 1 (related to figure 1). CosMx SMI chemistry and assay workflow.**

(a) Overview of CosMx SMI assay workflow. Fresh frozen tissue sections were hybridized with probes against RNA targets and protein segmentation markers, allowing simultaneous visualization of spatially restricted pathological features and RNA profiling at subcellular resolution (created with BioRender.com).

(b) Schematics of SMI ISH reporter design. The ISH reporter probe consists of two domains: a target recognition domain complimentary to the RNA target and a readout domain recognizable by reporter probes. The readout domain has four landing regions; each can be bound by a unique fluorescent reporter.

(c) Schematics of GeoMx SMI readout process. The sample goes through 16 cycles of reporter hybridization and quenching, generating a complete, unique color barcode for each gene *in situ*.

(d) Summary of the 950 plex RNA panel for this study. Many genes fall into multiple categories.

**Supplementary table 1. Quality control metrics of CosMx run.**

**Supplementary figure 2 (related to figure 2). Supervised cell type classification via negative binomial data generation model using the reference profile.**

(a) Reference gene expression profiles used in supervised cell type classification. This reference profile was based on a comprehensive cell typing census of mouse brain by single-cell RNA sequencing. The x-axis listed marker genes used in this profile. The y-axis listed the abbreviations and full names of each cell type.

(b) UMAP projection and clustering of cells based on the reference profile. 43 cell types have been identified.

(c) Condensation of cell types. To simplify downstream analysis, the cell types identified by the supervised algorithm (x-axis) are collapsed into higher levels (y-axis).

**Supplementary figure 3 (related to figure 4). The effect of tau pathology on thalamus and hypothalamus.**

(a) Spatial expression pattern of *Camk2a*. This gene shares the promoter driving the expression of mutant human tau transgene in rTg4510 brains.

(b) Body (left) and brain (right) weight of rTg4510 and WT mice (n = 4 per genotype). Significance was determined using unpaired t test; error bar represents SEM.

**Supplementary figure 4 (related to figure 5). Identification of spatial interaction domains.**

(a) Mapping of 9 spatial niches on the WT brain section. A single-cell neighborhood matrix was first computed to specify the number of each cell type among each cell’s closest neighbors within an 80 µm radius in 2D physical space. The resulting matrix for each tissue section was processed independently and clustered using the Mclust algorithm to identify spatial interaction domains. The description names of each domain were assigned by a neurologist.

(b) Representative spatial maps of niche 1 (cortex) in rTg4510 and WT brains. In WT brains, niche 1 captured cells from both the cortex and hippocampus. In rTg4510 brains, niche1 showed only partial co-localization with cortical areas, while the hippocampus was identified as a distinct niche, niche 2

**Supplementary figure 5 (related to figure 5). Identification of tangle bearing cells.**

(a) Pie chart showing the composition of cell types within 0 to 10μm of AT8-positive stains. Cell types were classified using a supervised cell type classification algorithm. Only cells from spatial niches 1 and 2 were included in the analysis. Percentages were calculated relative to the total number of cells within the defined 10μm radius of AT8-positive stains in rTg4510 brains and reported as the average value between the 2 biological replicates.

(b) Representative image showing the cell identities in close proximity to AT8-positive stains. AT8 staining is shown in yellow, with cell borders outlined in blue. White arrows indicate AT8- positive stains detected by the Fiji algorithm. Black asterisks denote cells located within 10μm of these stains. Colored dots represent specific cell types: red for neurons, green for oligodendrocytes, blue for astrocytes, and purple for microglia.

(c) Representative image showing the cell identities near false-positive AT8 stains. Similar to panel (b), AT8 staining is in yellow, with cell borders outlined in blue. White arrows indicate false-positive AT8 stains detected by the Fiji algorithm, while black asterisks mark cells within 10μm of these false positives. Colored dots indicate cell types, with orange representing vascular cells.

**Supplementary figure 6 (related to figure 6). Cross-validation between unsupervised (y-axis) and supervised clustering (x-axis).** Heat map showing the number of cells that fell within different cell types. To identify different OL lineages (red arrows), supervised cell typing was not condensed into higher level.

**Supplementary Table 2 (related to figure 6). Information of human samples.**

