## Supplemental table and figures for "Single-cell Spatial Transcriptional Profiling Uncovers Heterogeneous Cellular Responses to Pathogenic Tau in a Mouse Model of Neurodegeneration"

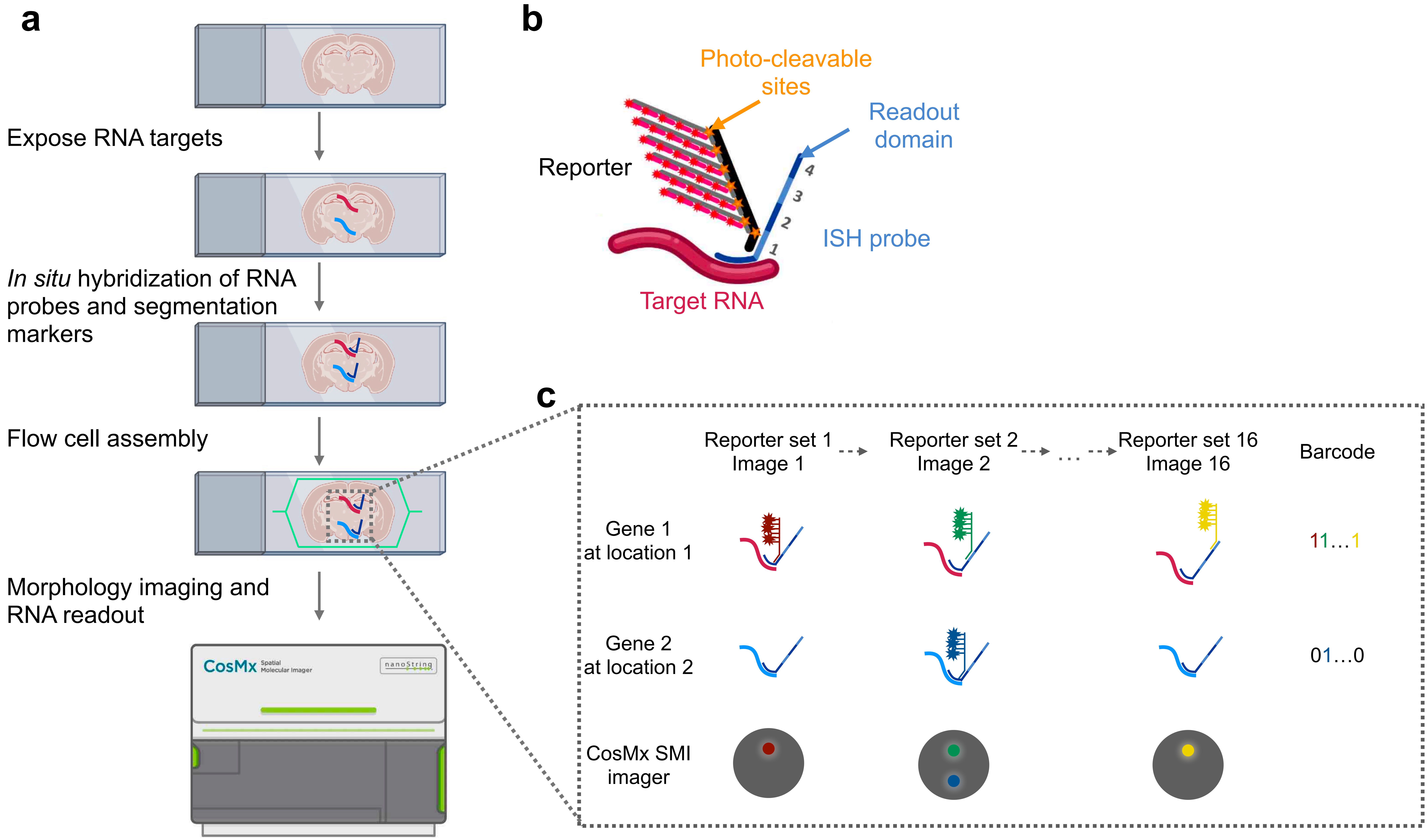


**d**

| Applications | # of genes |
|---|---|
| Cell typing and mapping | 215 |
| Glial activity | 226 |
| Neuropathology | 371 |
| Inflammation | 268 |
| Neurotransmission | 829 |
| Cell stress and damage response | 252 |
| Regulatory signaling | 635 |
| Extracelluar matrix reorganization | 57 |

Figure S1

| | slide_ID | R4775S1_rTg4510 | R4775S2_WT | R4784S1_rTg4510 | R4784S2_WT | all_slides |
|---|---|---|---|---|---|---|
| | Mouse_ID | m14213 | m14251 | m14227 | m14230 | NA |
| Basic info | FOV # | 68 | 95 | 72 | 81 | 316 |
| | cell # | 43987 | 70707 | 72412 | 79966 | 267072 |
| | % cells passing QC | 99.72 | 99.85 | 98.91 | 98.96 | 99.31 |
| | detected gene # | 725 | 788 | 804 | 809 | 788 |
| | % detected genes | 76.32 | 82.95 | 84.63 | 85.16 | 82.98 |
| Segmentation info | total transcript # | 2.36E+07 | 8.13E+07 | 4.21E+07 | 5.35E+07 | 2.01E+08 |
| | extracellular transcript # | 1.05E+07 | 3.91E+07 | 1.86E+07 | 2.40E+07 | 9.23E+07 |
| | % transcript assigned to cells | 55.48 | 51.84 | 55.81 | 55.15 | 54.51 |
| Volume info (5um thick) | volume measured (um$^3$) | 2.2E+08 | 3.07E+08 | 2.33E+08 | 2.62E+08 | 1.02E+09 |
| | transcript #/um3 | 0.107 | 0.265 | 0.181 | 0.204 | 0.198 |
| | intracellular volume (um$^3$) | 2.94E+07 | 5.92E+07 | 3.59E+07 | 4.32E+07 | 1.68E+08 |
| | intracellular transcript #/um$^3$ | 0.445 | 0.712 | 0.655 | 0.683 | 0.644 |
| count per cell | transcript #/cell | 297 | 596 | 325 | 369 | 405 |
| | Q1 of transcript #/cell | 90 | 181 | 74 | 77 | 106 |
| | Q9 of transcript #/cell | 571 | 1151 | 677 | 779 | 816 |
| | NegPrb #/cell | 0.173 | 0.319 | 0.147 | 0.106 | 0.185 |
| count per cell per target | transcript #/cell / plx | 0.313 | 0.627 | 0.342 | 0.388 | 0.426 |
| | NegPrb #/cell / plx | 0.0173 | 0.0319 | 0.0147 | 0.0106 | 0.0185 |

Table S1

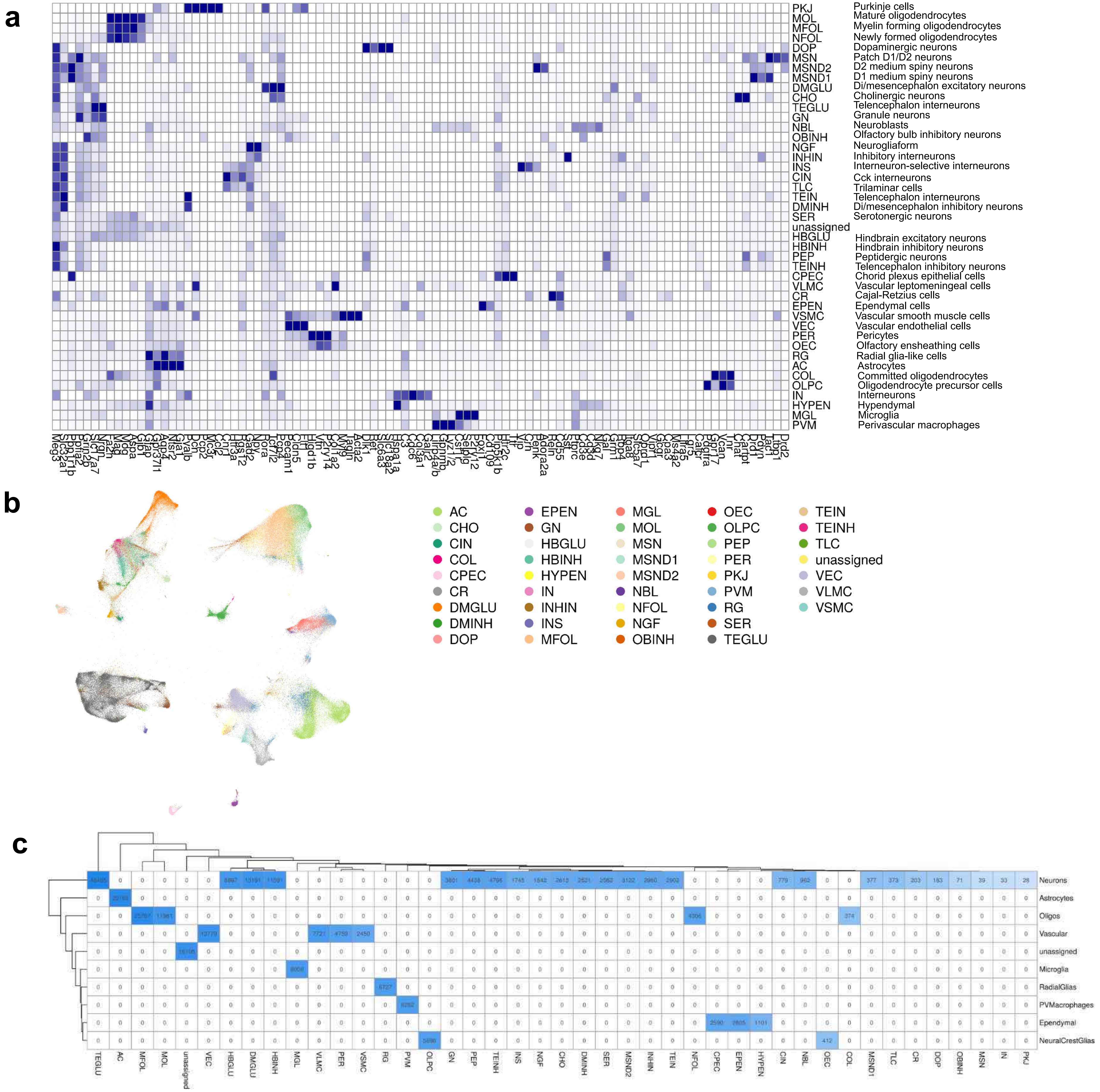
a
PKJ Purkinje cells
MOL Mature oligodendrocytes
MFOL Myelin forming oligodendrocytes
NFOL Newly formed oligodendrocytes
DOP Dopaminergic neurons
MSN Patch D1/D2 neurons
MSND2 D2 medium spiny neurons
MSND1 D1 medium spiny neurons
DMGLU Di/mesencephalon excitatory neurons
CHO Cholinergic neurons
TEGLU Telencephalon interneurons
GN Granule neurons
NBL Neuroblasts
OBINH Olfactory bulb inhibitory neurons
NGF Neurogliaform
INHIN Inhibitory interneurons
INS Interneuron-selective interneurons
CIN Cck interneurons
TLC Trilaminar cells
TEIN Telencephalon interneurons
DMINH Di/mesencephalon inhibitory neurons
SER Serotonergic neurons
unassigned
HBGLU Hindbrain excitatory neurons
HBINH Hindbrain inhibitory neurons
PEP Peptidergic neurons
TEINH Telencephalon inhibitory neurons
CPEC Chorid plexus epithelial cells
VLMC Vascular leptomeningeal cells
CR Cajal-Retzius cells
EPEN Ependymal cells
VSMC Vascular smooth muscle cells
VEC Vascular endothelial cells
PER Pericytes
OEC Olfactory ensheathing cells
RG Radial glia-like cells
AC Astrocytes
COL Committed oligodendrocytes
OLPC Oligodendrocyte precursor cells
IN Interneurons
HYPEN Hypendymal
MGL Microglia
PVM Perivascular macrophages
b
AC
CHO
CIN
COL
CPEC
CR
DMGLU
DMINH
DOP
EPEN
GN
HBGLU
HBINH
HYPEN
IN
INHIN
INS
MFOL
MGL
MOL
MSN
MSND1
MSND2
NBL
NFOL
NGF
OBINH
OEC
OLPC
PEP
PER
PKJ
PVM
RG
SER
TEGLU
TEIN
TEINH
TLC
unassigned
VEC
VLMC
VSMC
c
Neurons
Astrocytes
Oligos
Vascular
unassigned
Microglia
RadialGlias
PVMacrophages
Ependymal
NeuralCrestGlias

Figure S2

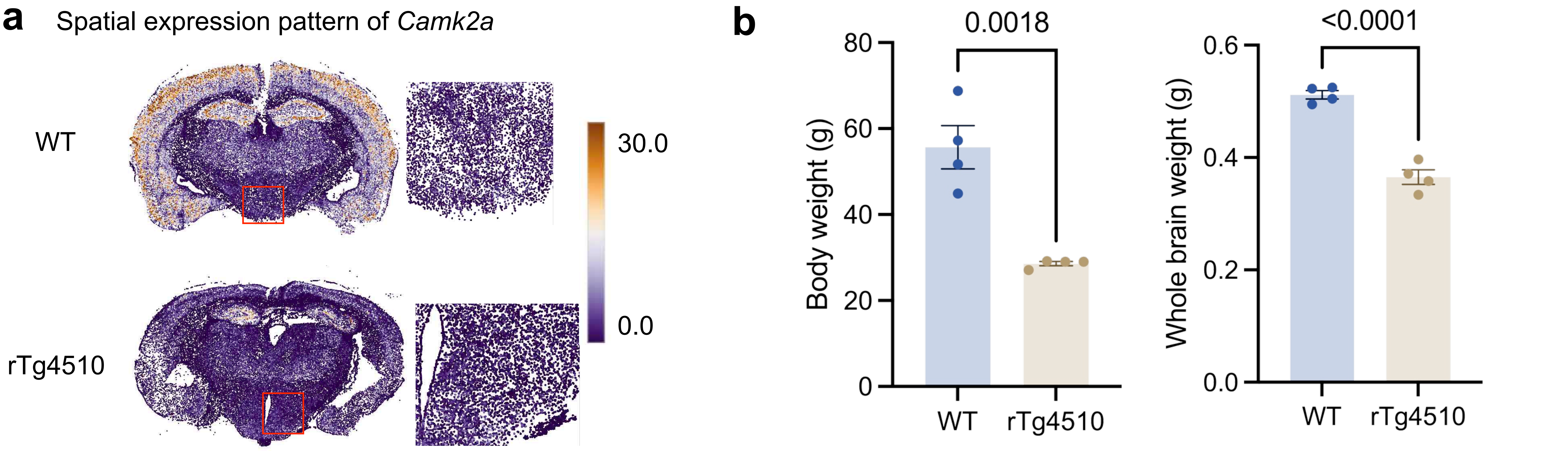
a
Spatial expression pattern of *Camk2a*
WT
rTg4510
30.0
0.0
b
Body weight (g)
0.0018
80
60
40
20
0
WT
rTg4510
Whole brain weight (g)
<0.0001
0.6
0.4
0.2
0.0
WT
rTg4510

Figure S3

a

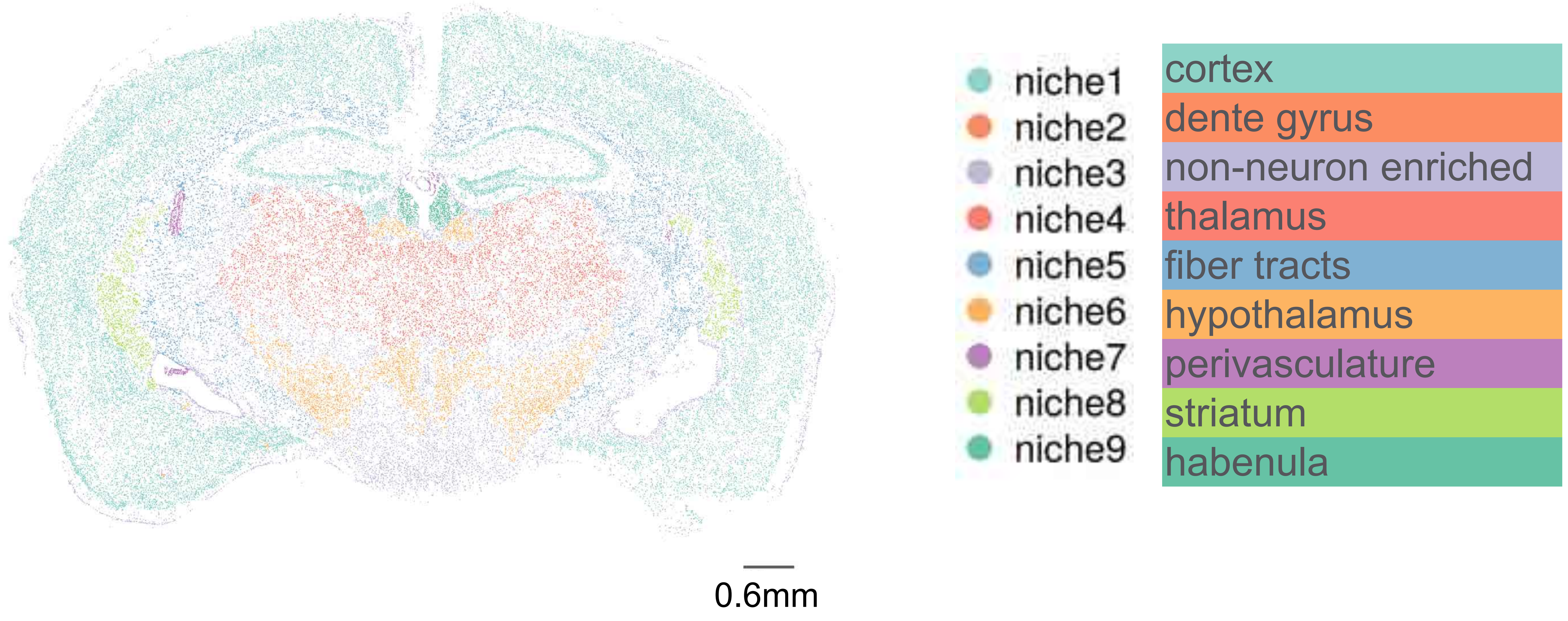
niche1
niche2
niche3
niche4
niche5
niche6
niche7
niche8
niche9
cortex
dente gyrus
non-neuron enriched
thalamus
fiber tracts
hypothalamus
perivasculature
striatum
habenula
0.6mm

b

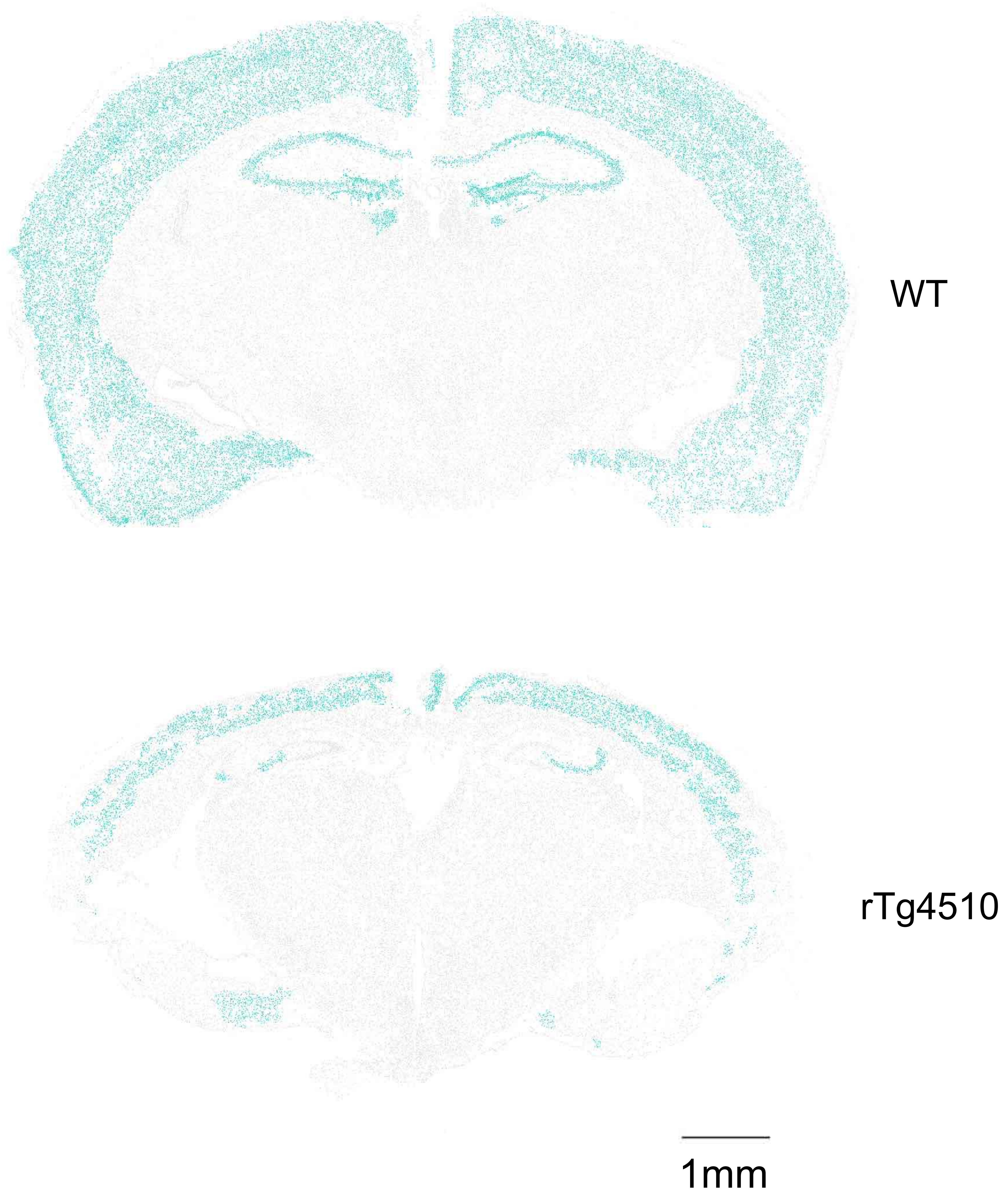
WT
rTg4510
1mm

Figure S4

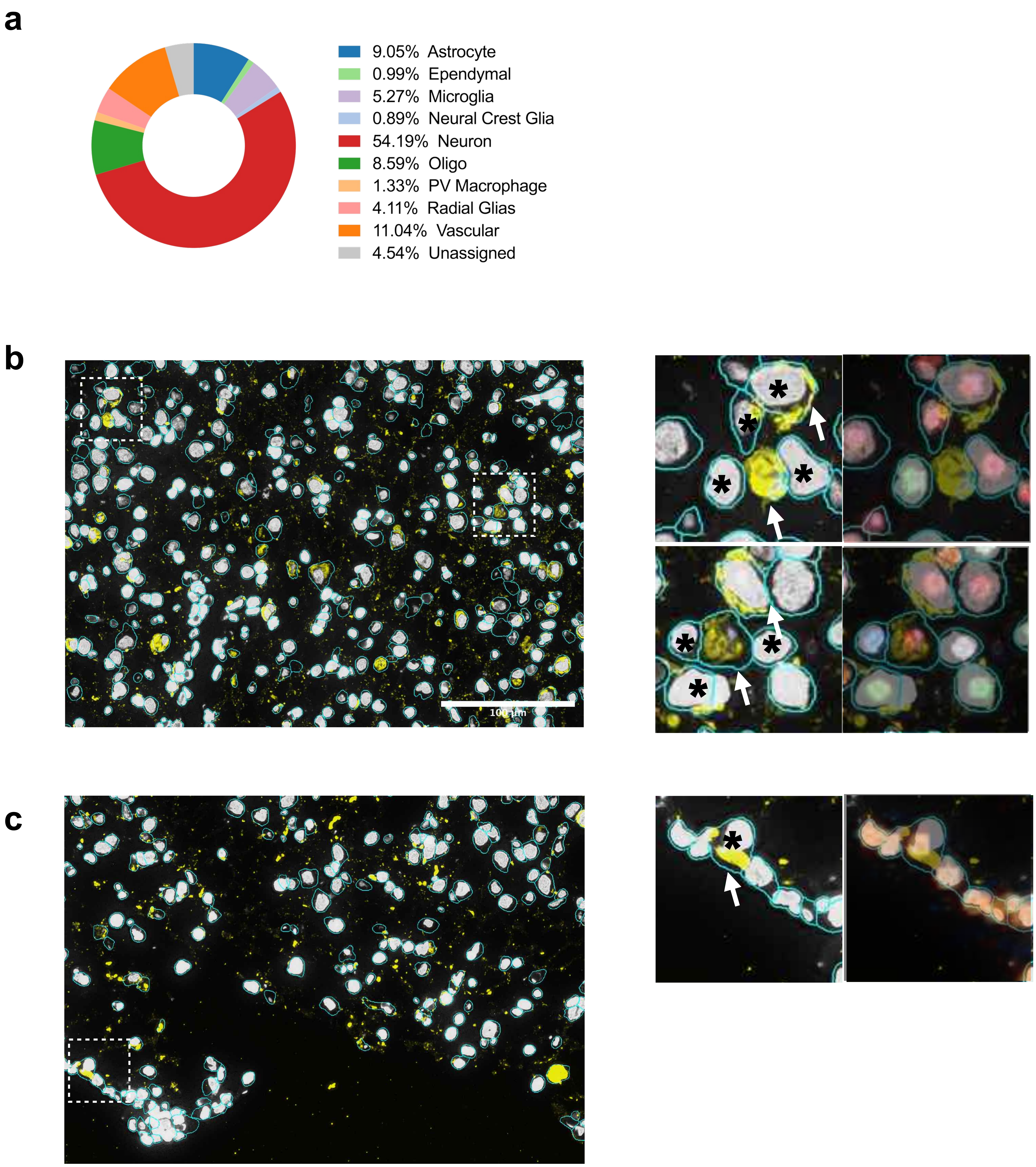
a
9.05% Astrocyte
0.99% Ependymal
5.27% Microglia
0.89% Neural Crest Glia
54.19% Neuron
8.59% Oligo
1.33% PV Macrophage
4.11% Radial Glias
11.04% Vascular
4.54% Unassigned
b
100 µm
c

Figure S5

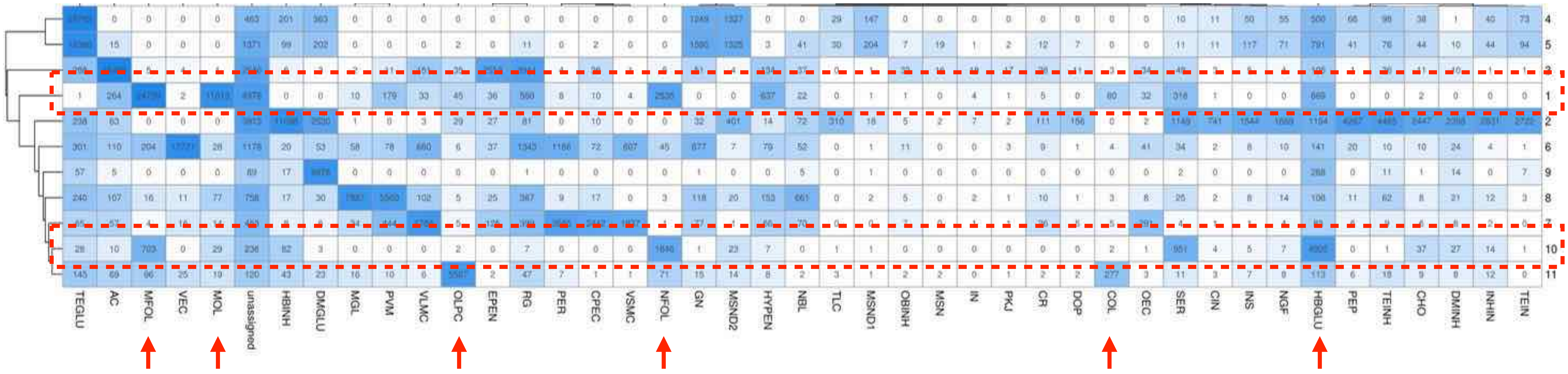
TEGLU
AC
MFOL
VEC
MOL
unassigned
HBINH
DMGLU
MGL
PVM
VLMC
OLPC
EPEN
RG
PER
CPEC
VSMC
NFOL
GN
MSND2
HYPEN
NBL
TLC
MSND1
OBINH
MSN
IN
PKJ
CR
DOP
COL
OEC
SER
CIN
INS
NGF
HBGLU
PEP
TEINH
CHO
DMINH
INHIN
TEIN
4
5
3
1
2
6
9
8
7
10
11

Figure S6

| | ID | Age | Gender | PMI | Amyloid | Braak | TangleTotal |
|---|---|---|---|---|---|---|---|
| | | | | PSP | | | |
| | 02-30 | 79 | M | 4 | B | II | 2.5 |
| | 02-34 | 82 | F | 1.83 | 0 | IV | 7.25 |
| | 04-59 | 93 | M | 2.83 | B | III | 3.25 |
| | 05-38 | 80 | M | 2.66 | 0 | III | 8 |
| | 09-23 | 99 | M | 4.33 | B | IV | 6 |
| | 09-24 | 90 | M | 2.5 | A | IV | 9 |
| | 10-01 | 98 | F | 2.5 | A | IV | 7 |
| | 11-69 | 73 | F | 2.71 | B | V | 9 |
| | 12-59 | 63 | M | 2.55 | 0 | V | 7.5 |
| | 12-60 | 64 | F | 2.63 | 0 | V | 11 |
| | 13-33 | 83 | M | 4 | 0 | V | 11 |
| | 14-35 | 90 | M | 3.5 | 0 | IV | 7.5 |
| | 15-12 | 84 | M | 5.57 | C | V | 9 |
| | 15-59 | 96 | F | 5.75 | C | V | 9 |
| Mean value | | 83.9 | N/A | 3.4 | N/A | N/A | 7.64 |
| | | | | ND | | | |
| | 13-40 | 73 | M | 4.12 | 0 | II | 2.25 |
| | 13-49 | 75 | F | 2.5 | 0 | II | 2.5 |
| | 14-19 | 90 | F | 2.5 | 0 | IV | 6.5 |
| | 14-42 | 92 | M | 2.3 | C | III | 5 |
| | 15-46 | 82 | F | 2.07 | A | I | 1.5 |
| | 15-60 | 82 | M | 3.33 | 0 | II | 3 |
| | 15-78 | 93 | F | 2.92 | B | III | 4 |
| | 16-11 | 90 | M | 5.42 | 0 | III | 6 |
| | 16-30 | 101 | M | 2.77 | C | IV | 8.5 |
| | 16-32 | 79 | M | 4.3 | 0 | II | 1 |
| Mean value | | 85.7 | N/A | 3.223 | N/A | N/A | 4.03 |

Table S2
